# On the enigmatic null effect of global methylation perturbation

**DOI:** 10.1101/2024.07.03.601377

**Authors:** Yifat Edrei, Asaf Hellman

## Abstract

Despite the established role of DNA methylation in transcriptional silencing, most of the genes remain unchanged upon its global erasing from the genome, while expression of other genes is actually raised. Since its discovery a few decades ago, this unsolved enigma has hampered true understanding of DNA methylation role. By applying an updated mathematical model to the results of the original studies, we showed that this paradoxical effect is due to between-gene differences in the balance between methylation-guided enhancers and silencers, which protect particular disease-causing genes from the otherwise harsh effects of global methylation perturbation. Nevertheless, local methylation alternation may still affect these genes. The results a settle a long-standing wondering and suggest that DNA-methylation regulatory role is significantly greater than previously thought.

## Main text

Global perturbation of the DNA methylation blueprint is a pan-cancer phenomenon ^1,2^. DNA methylation mediates transcriptional silencing by restraining the binding of transcriptional activators, and/or attracting transcriptional suppressors, to regulatory DNA targets ^3^. However, in spite of this established silencing effect, induction of genome-wide hyper or hypo-methylation typically leaves the most of the genes unaffected, whereas considerable fractions of the other genes altered in the opposite way to the expectation, i.e., increasing expression when methylation is globally declined, or vice versa. These unexplained inconsistencies have challenged the understanding of the role of DNA methylation in normal and transformed cells.

Whereas promoter methylation is negatively associated with gene expression level, both positive and negative associations between methylation and expression levels were reported among distal regulatory methylation sites ^4-9^. We recently demonstrated that expression level of cancer genes may be described as the sum effect of multiple, methylation-related positive and negative effects on transcription^10^. Considering these findings, we wondered whether the balance between positive and negative sites influences the effect of global methylation perturbation on particular genes.

We explored this hypothesis by mapping cis-regulatory methylation sites that were associated with the expression level of genes across human cancer genomes. Applying the algorithm^10^ to an open dataset of DNA methylation and gene expression levels in 280 colon cancers (Figure 1A), we located 30,886 gene-associated methylation sites, of which 18,025 showed negative association with gene expression, and 12,861 showed positive association (Figure 1B and Table S1). Both positive and negative sites carried the chromatin variant H3 lysine 4 mono-methylation (H3K4me1), hallmark of distal regulatory elements, in colon cells (Figure 1C) as well as in glioblastoma genomes ^10^. Of 6,023 associated genes, 3,910 (64.9%) were associated with more than one site (Figure 1D and Table S2).

**Figure 1.**
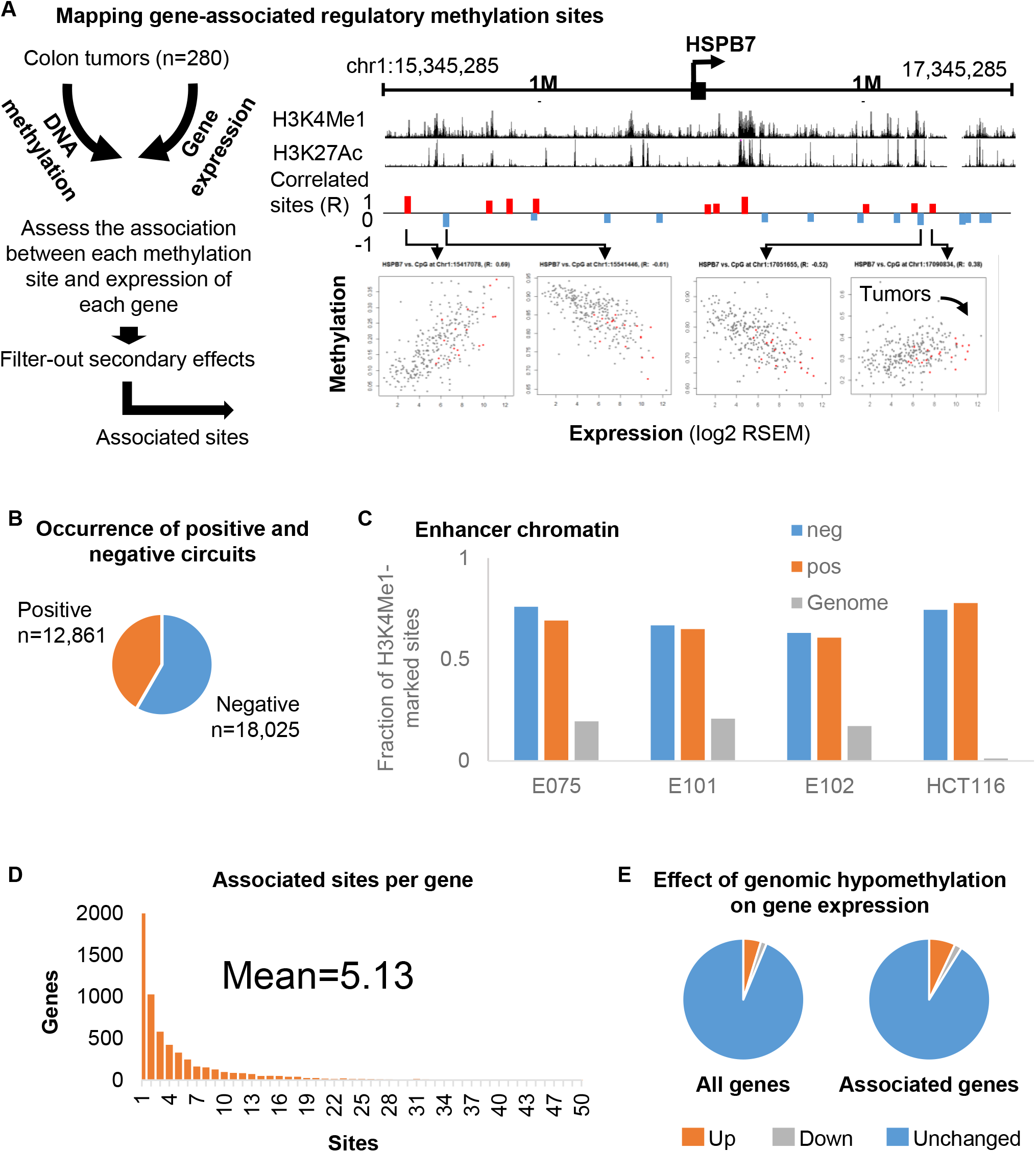
Cis-residing regulatory methylation sites associated with gene expression levels. **A**. Methylation sites within 2 mega base-pair intervals around gene transcription start sites were tested for association with the expression level of the related genes across colon cancers. **Left:** Scheme of the analysis. **Right:** Example of a mapped region around the HSPB7 gene. Genomic locations, colon chromatin marks of enhancer activity, and associated positive (red) or negative (blue) methylation sites are shown. Representative scatter plots of methylation versus expression levels across 280 colon cancers are shown in the lower panel. Scattered red dotes denote normal (untransformed) colon samples. **B**. Occurrence of methylation sites showing positive or negative association with gene expression. **C**. Fractions of positive or negative methylation sites carrying the histone-3-lysine-4-mono-methylation marker of enhancer chromatin in various colon cell lines. Genomic frequencies if this marker are given for reference. **D**. Distribution of the number of methylation sites per associated gene. **E**. Fraction of genes that gained, reduced, or unaltered expression levels upon induced global genomic hypomethylation. **Left:** All analysed genes. **Right:** genes with associated methylation sites.

To assess the effect of methylation mutation on the associated genes, we re-analyzed gene expression data^11,12^ of colon cancer cells lacking the enzymes that establish and maintain DNA methylation patterns. In these cells, the global DNA methylation level is diminished by more than 80 percent relative to the untreated cells, throughout the genome ^13,14^. Despite this global de-methylation, only 4.7% of the genes (n=617) showed increased expression levels as compared to the wild type cells. Moreover, a two-fold or greater reduction in expression levels was observed among 1.6% (n=205) of the genes. The remaining 93.7% (12,250) of the genes showed less than two-fold transcriptional difference upon global demethylation, as compared to the wild type cells. Similar patterns of gained, decreased, and unaltered expression levels observed in the subgroup of genes for which we located associated methylation sites (Figure 1E).

Next, we explored the cis-regulatory structures of genes with multiple associated regulatory-sites. To this end, redundancy versus synergic interactions between the associated sites were evaluated by comparing the effect on expression of particular sites versus the combined effect of all associated sites. For example, the CHMP7 gene associated with six negative and sixteen positive sites, of which the maximum correlation values was 0.495. A cumulative model considering all sites provided improved prediction (R=0.779), indicating regulatory synergism among the sites (Figure 2A and Table S2). Of the 3,910 genes with multiple associated sites, 3,494 (88.2%) were controlled by networks of synergic cis-regulatory sites (Figure 2B and Table S2). Moreover, 1,937 (56.2%) of these networks comprised both positive and negative sites (Figure 2C and Table S2). We then asked whether the balance between positive and negative regulatory sites predicts the effect of global methylation perturbation. Strikingly, genes that showed increased expression upon demethylation, were associated with more negative than positive regulatory sites, and genes that showed decreased expression were associated with more positive than negative sites (p=0.013) (Figure 2D). Hence, global de-methylation gains the expression of genes that associated with excess of negative sites, and reduces the expression of genes with more positive sites.

**Figure 2.**
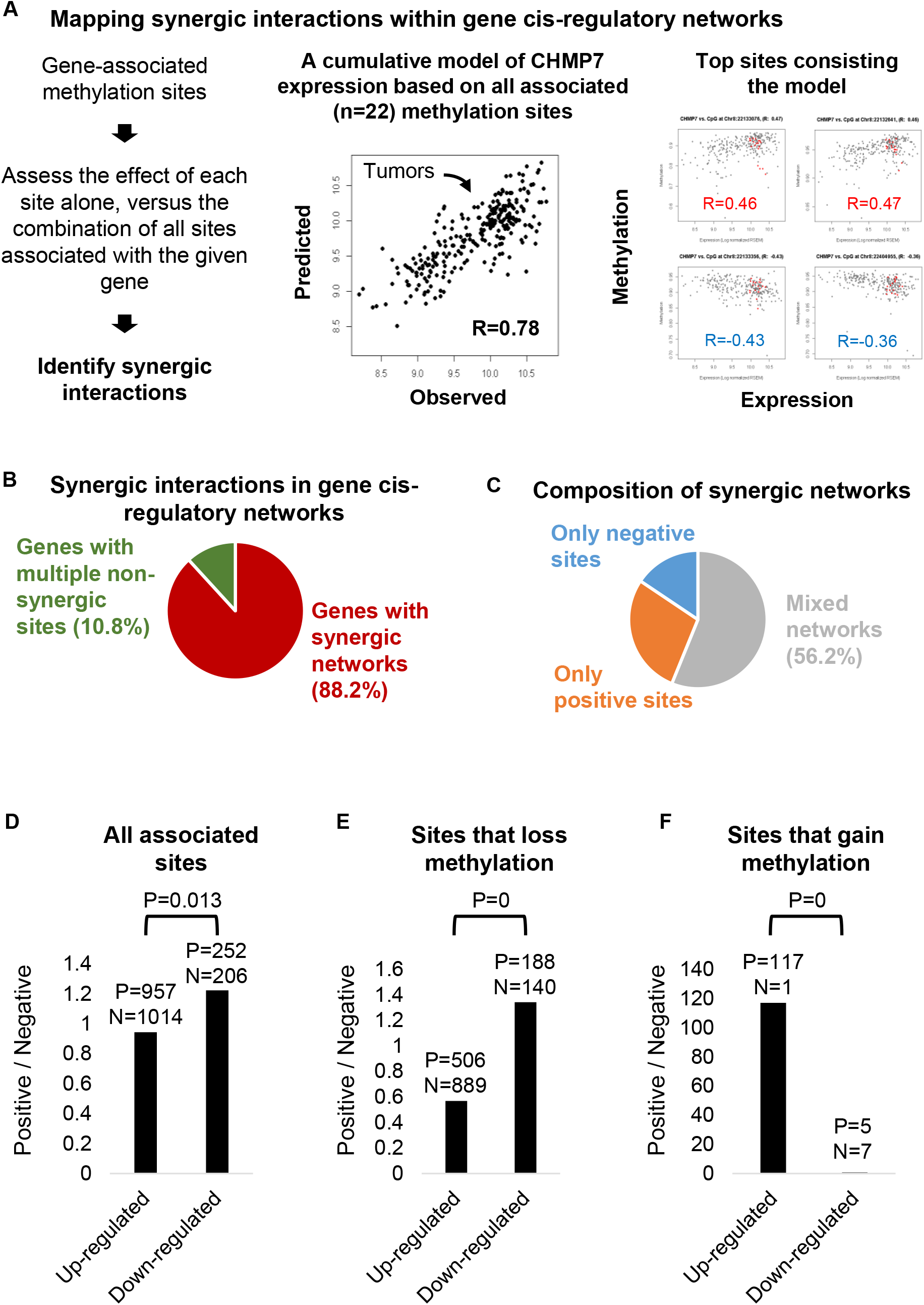
Balance between positive and negative methylation sites defines the effect of global methylation perturbation. **A**. Analysis of synergic interactions between gene-associated methylation sites. **Left:** Scheme of the analysis: Effects of discrete cis-residing methylation sites on gene expression across the tumors, were compared with combinatorial models considering all the methylation sites associated with the given gene. Synergic models were identified as those convening stronger effect on expression than each of the associated sites alone. **Right:** Methylation-based prediction of CHMP7 expression. The model includes 22 positive and negative associated methylation sites providing correlation (R) values between 0.475 to - 0.495. The cumulative model provided a supreme effect on expression as compared to each of the sites alone, indicating synergic interactions between at least some of the sites. **B**. Fraction of gene cis-regulatory networks showing synergic interactions. **C**. Fractions of synergic networks that incorporated only negative, only positive, or both negative and positive methylation sites. **D**. Ratios between the number of positive and negative sites that were associated with genes that responded with increase or decrease expression upon global hypomethylation. Actual numbers of associated positive and negative sites shown for the gene groups. **E**. Same as D, but for sites that decreased their methylation level by at least 10% upon induced global hypomethylation. **F**. Same as D, but for sites that increase their methylation level by at least 10% upon induced global hypomethylation.

We also noted that some sites maintained high methylation levels despite the general hypomethylation of the manipulated cells, possibly due to remaining methylation marks and/or residual re-methylation activity. Repeating the analysis among the sites that actually showed methylation loss (at least 10% methylation reduction), revealed an even more prominent tendency of genes with excess of negative sites to increase expression levels, and of genes with a more positive sites to decrease expression (p=0) (Figure 2E). Furthermore, among the small minority of sites that actually gained methylation in the manipulated cells, increased gene expression was predominantly supported by positive methylation sites, and expression loss by negative methylation sites (p=0) (Figure 2F). Thus, genes were responded by elevating or lowering expression levels, to methylation gain or loss, depending on the nature of methylation sites comprising their cis-regulatory networks.

In conclusion, we showed that gene misregulation is dictated by the particular structure of gene cis-regulatory networks, modulated by the methylation alterations of regulatory sites. Our results show that regulatory schemes incorporating both positive and negative sites are frequent in the human genome (Figure 2C) ^10^, and that the effect of global methylation perturbation is modified by the contradicting regulatory elements composing them (Figure 2D-F). This unique organization of cis-regulatory networks may serve as a barrier against acute loss of control over the cellular transcriptional program in case of main epi-patterning failures. However, while protecting from genome-wide hyper or hypo methylation, this mechanism allows site-specific methylation mutations to be highly effective. As we elsewhere showed^10^, such differential methylation effects are frequent in cancer genomes, and may explain a significant portion of inter-individual gene-expression variation.

Our findings also shed light on the relatively negligible effect of global methylation perturbation, formerly perceived as indication for a limited role of methylation mutation in transformed cells. Through systematic assessment of cis-regulatory networks, the effect of global or local methylation mutations may be better evaluated, and be taken into account when prognosis of particular epi-mutations is made, or particular epi-therapeutic protocols are considered.

