## supplementary materials for "On the enigmatic null effect of global methylation perturbation"

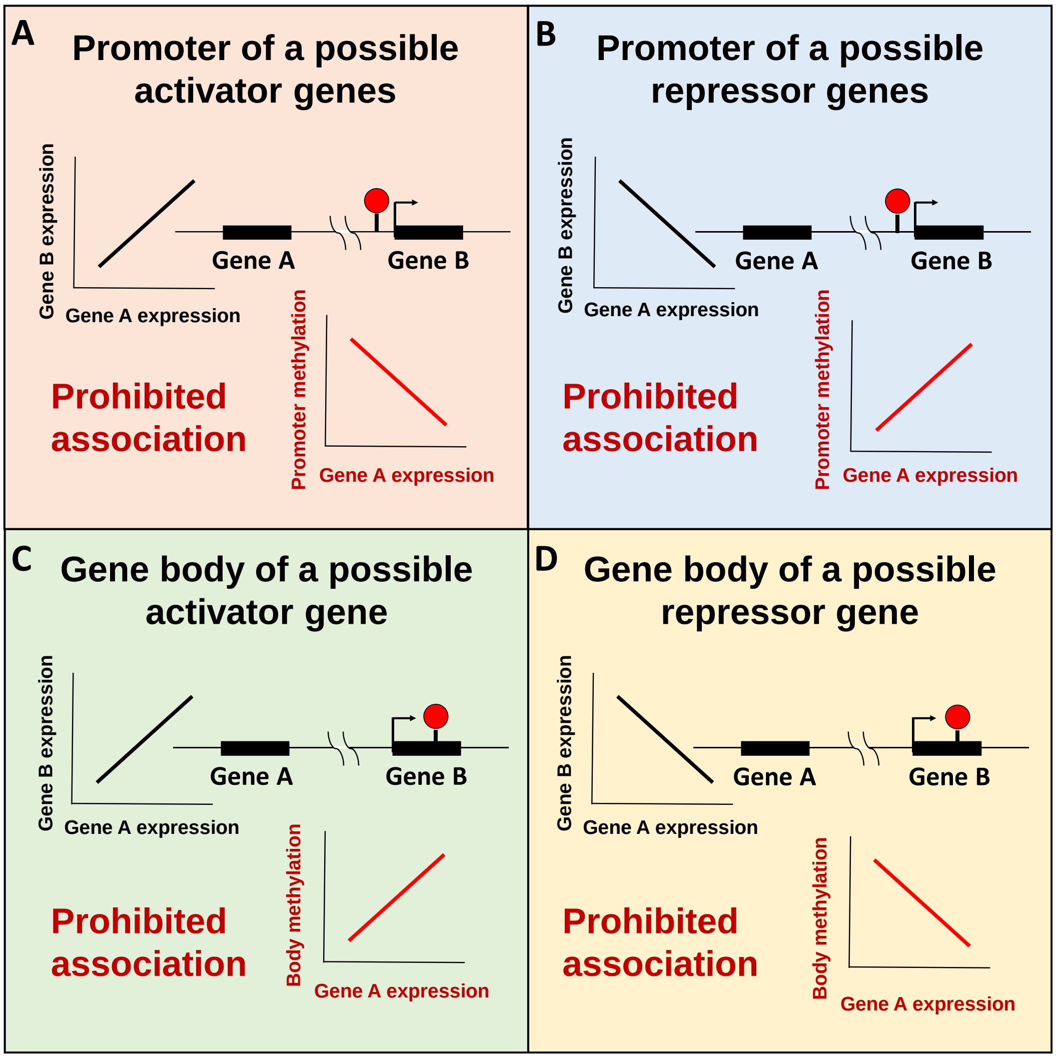


Figure S1. **Eliminated associations due to possible secondary effects. A.** Prohibited association between methylation of a promoter site and expression of a possible activator of the indicated gene A. **B.** Prohibited association between methylation of a promoter site and expression of a possible repressor of the indicated gene A. **C.** Prohibited association between methylation of a gene-body site and expression of a possible activator of the indicated gene A. **D.** Prohibited association between methylation of a gene-body site and expression of a possible repressor of the indicated gene A.

**Supplementary methods**

Level 3 Illumina HumanMethylation450 BeadChip methylation data and level 3 RNAseq (RNAseqV2 normalized RSEM) expression data of colorectal adenocarcinomas (COAD, n=280) were obtained from the Cancer Genome Atlas (TCGA) (https://tcgadata.nci.nih.gov/tcga/dataAccessMatrix.html). H3K4me1 enhancer chromatin data of digestive cell lines were downloaded from the ENCODE (wgEncodeSydhHistoneHct116H3k04me1UcdPk.narrowPeak.gz) and the Roadmap (E075-H3K4me1.broadPeak.gz, E101-H3K4me1.broadPeak.gz, E102-H3K4me1.broadPeak.gz) public databases. Expression data (RNAseq) of HCT-116 colon cancer (HCT116WT) cells, and HCT-116 DNMT1 and DNMT3B double methyltransferase-knockdown (HCT116DKO) cells [11,12], were downloaded from the Gene Expression Omnibus (GEO) database accession numbers GSE39068 and GSE60106. BeadChip methylation data for HCT116WT and HCT116DKO were downloaded from the GEO accession number GSE29290. Gene-associated methylation sites were mapped by applying pairwise Spearman's rank correlation coefficient with Bonferroni correction for multiple-hypothesis testing. Sites that also displayed a prominent ($R^2$ > 0.1) correlation with expression of the pan-blood cells marker CD45 were considered a possible result of blood contamination of the tumor sample and were eliminated from the analyses, as described [15]. Potential secondary effects were considered in cases were the associated site was included within the transcribed portion (the gene body, excluding the first 5kbp) of another gene, or located within the promoter (-1500 to +2500 bp of TSS) of another gene. For these cases, a correlation between the expression level of the associate and the hosting genes was assessed, and interactions with $R^2$>0.1 that fit one of the scenarios described in Figure S1, were excluded from the analysis. Expression prediction models were calculated by applying multiple linear regression. Model p-values were evaluated by ANOVA for Linear Model Fit. Models with p-value < 0.05 were considered significant. p-values of the differences between the number of positive and negative methylation sites associated with up-regulated or down-regulated genes were analyzed by 2X2 chi-squared test at a significance level <0.05.
